# Analysis of Lower-Grade Gliomas in MRI Through Segmentation and Genomic Cluster-Shape Feature Correlation

**DOI:** 10.1101/2022.08.01.502342

**Authors:** Rohit Paradkar, Ria Paradkar

**Affiliations:** Beth Israel Deaconess Medical Center; Albany Medical College

## Abstract

Gliomas, which originate from glial cells, are considered the most aggressive type of brain tumors. Currently, glioma research efforts are focused mainly on high-grade gliomas. This project aims to analyze lower-grade gliomas (LGG) in MRI and extend the understanding of LGGs. LGG segmentation, which outlines tumors in MRIs, is crucial to developing effective treatment plans. However, segmentation performed manually by radiologists is tedious, time-consuming, and often leads to inter-observer variability. Another unexplored area of LGG research is genomic subtypes. These subtypes can play a large factor in how LGGs can be treated, however there is currently no noninvasive method of identifying these subtypes. Recent studies suggest that LGG shape features have a correlation with genomic subtypes and should be investigated as a viable factor in LGG treatment options. This presents a need for additional research as most LGGs eventually develop into high-grade gliomas. The specific aims of this project include analyzing LGGs through deep learning-based segmentation, shape feature extraction, and statistical analysis to identify correlation between selected shape features and genomic subtypes. To realize these goals, programs were written and run using a publicly available LGG dataset. In terms of automatic segmentation, two models were created using different convolutional neural networks (CNN). The highest performing model used U-Net with a ResNeXt-50 backbone and yielded a 91.4% accuracy in terms of mean intersection over union (IoU) after testing. Shape feature extraction included three geometric features and 4 radiomic features which quantified tumor shape in 2D and 3D. Angular Standard Deviation (ASD), Margin Fluctuation (MF), Bounding Ellipsoid Volume Ratio (BEVR), Elongation, Major and Minor Axis Length, and Volume were tested for correlation with genomic subtype data using 49 Pearson’s chi-squared tests. P-values less than or equal to 0.05 suggested correlation. Statistical analysis found 16 statistically significant associations. The strongest associations were between MF and the RNASeq cluster (p < 0.00003), ASD and the RNASeq Cluster (p < 0.0005), and volume and the RPPA cluster (p < 0.0035).

## 1) Introduction & Background Information

Gliomas are brain tumors that originate from glial cells and are considered the most severe type of brain tumor. They are divided into 4 grades based on tumor malignancy, with grades II and III being considered lower-grade gliomas (LGG). [1] Because glioblastomas (Grade IV) are the most common form of malignant brain cancer and are considered one of the deadliest human cancers, there has been significant demand for high-grade glioma (HGG) research in the past. [2] Despite considerable emphasis on high-grade gliomas, LGG research funding is low due to its higher survival rate. This creates a major problem, as LGG is infiltrative and eventually develops into HGG if untreated. [3] This makes it necessary to increase research efforts relating to diagnosis and the process of treatment for lower-grade gliomas.

One of the first steps in diagnosis is computer imaging, which for LGG, is generally done through magnetic resonance imaging (MRI). MRI utilizes computers to generate detailed images of the brain. The next step is segmentation, which is the process of identifying and outlining the tumor region inside the MRI image of the brain. Currently, segmentation is done manually by neuroradiologists or clinicians. [4] Despite manual segmentation being done by experts and often being the most reliable method of segmentation, it is very time consuming and often leads to inter-observer variability, which can significantly sway how MRI images are interpreted by different professionals. [4] Because of these drawbacks, one effective solution is automatic LGG segmentation, as a predictive deep learning-based model would provide quick and consistent segmentation and yield substantial improvements to diagnosis and treatment. Past automatic LGG segmentation models have been effective; however, many do not consider both pre- and post-operative data. [6,7] Considering images with and without tumors will increase model flexibility and enable more efficient training.

In addition to segmentation, inter-observer variance can also affect the process of LGG treatment. One promising approach that can aid doctors in treating LGG is genomic subtypes. Several studies suggest that, using various sequencing and proteomic assessments, LGG can be associated with different genomic subtypes, which are important factors in determining the course of treatment. There is no noninvasive method of identifying these genomic subtypes. Previous literature indicates correlation between LGG shape characteristics and subtypes. [5,7,8] This creates a need to conduct radiogenomic analysis and strengthen inferences about these correlations. Past projects have conducted similar analysis; however, this project aims to retest previously identified shape features while also assessing potential correlation between genomic subtypes and additional shape features. [5,8] Previous papers have only extracted shape features coming from ground-truth tumor images rather than segmented tumor predictions. [5,7,8]

This project aims to create a pipeline which will 1) create a deep learning-based model that performs automatic segmentation of LGG in MRI, 2) extract radiomic and geometric shape features from predicted tumor regions, and 3) use statistical analysis to infer correlation between select shape features and genomic subtypes.

## 2) Materials & Data

### 2-1) Materials

- MacBook Laptop with Internet connection
- Anaconda Navigator
- Spyder IDE 4.1.5
- Python v. 3.8
- Jupyter Notebook
- Google Colaboratory with GPU
- Lower Grade Glioma (LGG) MRI Image dataset from The Cancer Imaging Archive (TCIA) and Genomic Subtype data from The Cancer Genome Atlas (TCGA)
- Python packages - pytorch, torchvision, matplotlib, numpy, pandas, albumentations, scikit image, scikit learn, opencv, glob, pyradiomics, scipy, seaborn, heatmapz, etc.

### 2-2) Data

The data utilized in this project had two sources: The Cancer Imaging Archive (TCIA) and The Cancer Genome Atlas (TCGA). [9,10] This data included information about 110 different patients from five different institutions diagnosed with LGG. The dataset included patients with varying histological grades, tumor location, gender, age at initial diagnosis, race, and ethnicity. Refer to Appendix to see data visualization images.

The imaging data originating from TCIA included both MRI images and T2-Weighted Fluid Attenuated Inversion Recovery (FLAIR) abnormality masks (image of tumor region outlined in white pixels on black background) for all patients. Each image had a corresponding mask, and all images were either T-1 or T-2 weighted. T-1 and T-2 are different MRI pulse sequences which have noticeable differences in image brightness and appearance. Masks were automatically segmented initially, and then manually annotated by radiologists. All 110 patients had multiple images, with each patient having a range of 20-90 images over the course of treatment.

Genomic data coming from TCGA correspond to the same 110 patient images included in the TCIA dataset. This collection includes information about genomic clusters occurring in patient images:

RNASeqCluster - This cluster includes data regarding findings from RNA sequencing, a form of next-generation sequencing. The values range between 1-4, with each value signifying one of the four different genome clusters.

Methylation Cluster - This cluster includes data regarding findings from DNA Methylation Profiling. The values in this cluster range between 1-5, with each value correlating to 5 different DNA methylation patterns occurring in the LGG dataset.

miRNA Cluster - This cluster includes data regarding findings from miRNA sequencing. The values range between 1-4, with each value signifying one of the four genome clusters found in sequencing.

CN Cluster - The data values included in the CN Cluster range from 1-3. The values in this cluster correspond to 3 different classes of DNA copy number occurring in the dataset.

RPPA Cluster - The RPPA cluster data was obtained from results of reverse-phase protein lysate array profiling (RPPA). The data includes values ranging from 1-3, providing information about protein expression profiles in LGG.

Oncosign Cluster - the Oncosign values range from 1-3 and provide tumor categorization based on similarities in recurrent mutations and copy number variance.

CoC Cluster - Cluster of Cluster data ranges from 1-3. It provides a second-level clustering of class assignments derived from each individual subtype platform.

## 3) Design & Methods

### 3-1) Performance Criteria

#### Automatic Segmentation

- Total model training time should not exceed 7200 seconds (2 hours)
- Mean training Dice Coefficient should be greater than or equal to 0.85
- Mean testing intersection over union (IoU) should be greater than or equal to 90%
- Mean loss should be less than or equal to 0.15
- Model cannot overfit or underfit during training; model testing accuracy must be consistent with model training accuracy
- Model is measurable after every epoch

#### Shape Feature Extraction

- Shape features must be assessable for all patient image data
- 2D imaging must be transformable to 3D to extract quantitative features

#### Shape Feature - Genomic Subtype Correlation

- Quantitative data must be transformable to categorical data for use in Pearson’s Chi-Squared Tests

### 3-2) Methods/Design

#### Deep Learning Based Automatic LGG Segmentation

The first step was writing the programs for data collection, data preparation, and data visualization which would be fed into the CNN. Following preliminary steps, the dataset was split by the model so that 15% was set aside for model testing. 76.5% of image data was used to train the deep learning model, and 8.5% was utilized for training validation. Data augmentation was used to artificially create more training data by applying image transformations to existing data. These transformations included resizing, rotation, brightness filters, and other image alterations. Because of U-Net’s segmentation capabilities, it was used as the CNN for the prototype model. [11] Different layers of the CNN were defined through pytorch implementation and data was fed into the model for training. Validation methods were used to give basic accuracy metrics for training using the dice similarity coefficient and the BCE Dice loss function. Several parameters were constrained to maximize training efficiency and accuracy. Batch size, learning rate, and number of epochs were adjusted to 0.0001, 16, and 50 respectively. After training was completed, the model created tumor region predictions based on test data. The metric for model accuracy was the mean IoU of test images. It was found by comparing segmentation predictions and ground truth FLAIR masks. To measure accuracy and account for outlier predictions, the model was run 5 times and the IoU was averaged. After prototype model testing was completed, further research was conducted to enhance accuracy. One encouraging prospect was to include a convolutional backbone in the U-Net architecture to supplement feature-extraction networks during training. [12] The model was redesigned by integrating the ResNeXt-50 backbone with the CNN architecture and then retested using optimized parameter constraints and the same model evaluation metrics.

#### Radiomic & Geometric Shape Feature Extraction

Following LGG segmentation, the next step was extracting shape features from segmented tumors. Seven quantitative shape features which had been noted by several publications to be of importance in LGG radiogenomics were extracted. [5,7,8] All 3D shape features required transformation from 2D to 3D imaging using the JoinSeries method of SimpleITK. Three of the shape features were mathematically computed and four were computed using the pyradiomics package from Python:

##### Angular Standard Deviation (ASD)

ASD is a metric which can help in defining the circularity of a 2D object. It was calculated by finding the centroid of the tumor and drawing 10 equiangular segments from the centroid to the perimeter. The mean length of the segments was found. Then the standard deviation was calculated, with a smaller standard deviation corresponding to higher tumor circularity. To account for the fact that standard deviations could be higher if a tumor is larger despite a similar level of circularity between small and large tumors, radial distances for all images were standardized by dividing individual distances by mean distance.

##### Margin Fluctuation (MF)

MF is a metric which defines the smoothness of a tumor’s perimeter by quantifying the number of fluctuations observed. To compute it, the centroid of tumor regions was found. Next, the average radial distance from the centroid to the tumor boundary was calculated. Filtering was then applied on the tumor’s outer boundary to smoothen the perimeter such that the smoothened perimeter length was equal to 10% of the original perimeter length. Radial distances from the centroid to the filtered perimeter were evaluated. Next, the average difference between original perimeter radial distances and smoothened perimeter radial distances was computed. The standard deviation of the average difference was finally found to determine tumor spiculation. Radial distances were standardized to eliminate value penalization for larger tumors.

##### Bounding Ellipsoid Volume Ratio (BEVR)

BEVR is a metric which defines the 3D regularity of a LGG tumor. It is the ratio of the tumor’s volume to the volume of the smallest ellipsoid enclosing the tumor. To calculate this, a program sorted patient images and only took the largest tumor cross section for each patient. Using the pyradiomics package, the tumor volumes were calculated. Next, to find the volume of bounding ellipsoids, all coordinate points on tumor surface areas were inputted into a function from a freely available GitHub repository which used the Khachiyan Algorithm. [14,15] Finally, the BEVR was computed as a ratio. BEVR values closer to 1 signify more tumor regularity.

##### Radiomic Features

The other 4 shape features extracted from tumor images were determined by using the pyradiomics package. These features were tumor elongation, major and minor axis lengths, and tumor volume.

#### Shape Feature - Genomic Subtype Analysis

The final part of the model consisted of testing the potential correlation between the previously listed genomic subtypes and shape features. To do this, statistical tests between each shape feature and each genomic subtype were performed. Because genomic data was only categorical and shape feature data was quantitative, shape features had to be categorized by quartiles. Programs were written which identified values for quartile cutoffs and shape feature data points for each patient were categorized in the range of 1-4. To analyze the relationship between the two categorical variables, Pearson’s chi squared tests were run between each shape feature and genomic subtype. 49 total tests were performed. P-values convey the probability that the null hypothesis is true, meaning that no correlation between variables exists. P-values lower than .05 were interpreted as suggestive of potential correlation (≥ 95% chance of alternative hypothesis that correlation exists). Chi-squared statistic values were also reported for effect size measure in addition to degrees of freedom.

## 4) Results

### Automatic Segmentation

After training and testing the U-Net segmentation model and then redesigning using the ResNeXt-50 convolutional backbone, accuracy of the prototype and redesigned models were calculated. Figure 1 compares the two using performance metrics.

**Figure 1.**
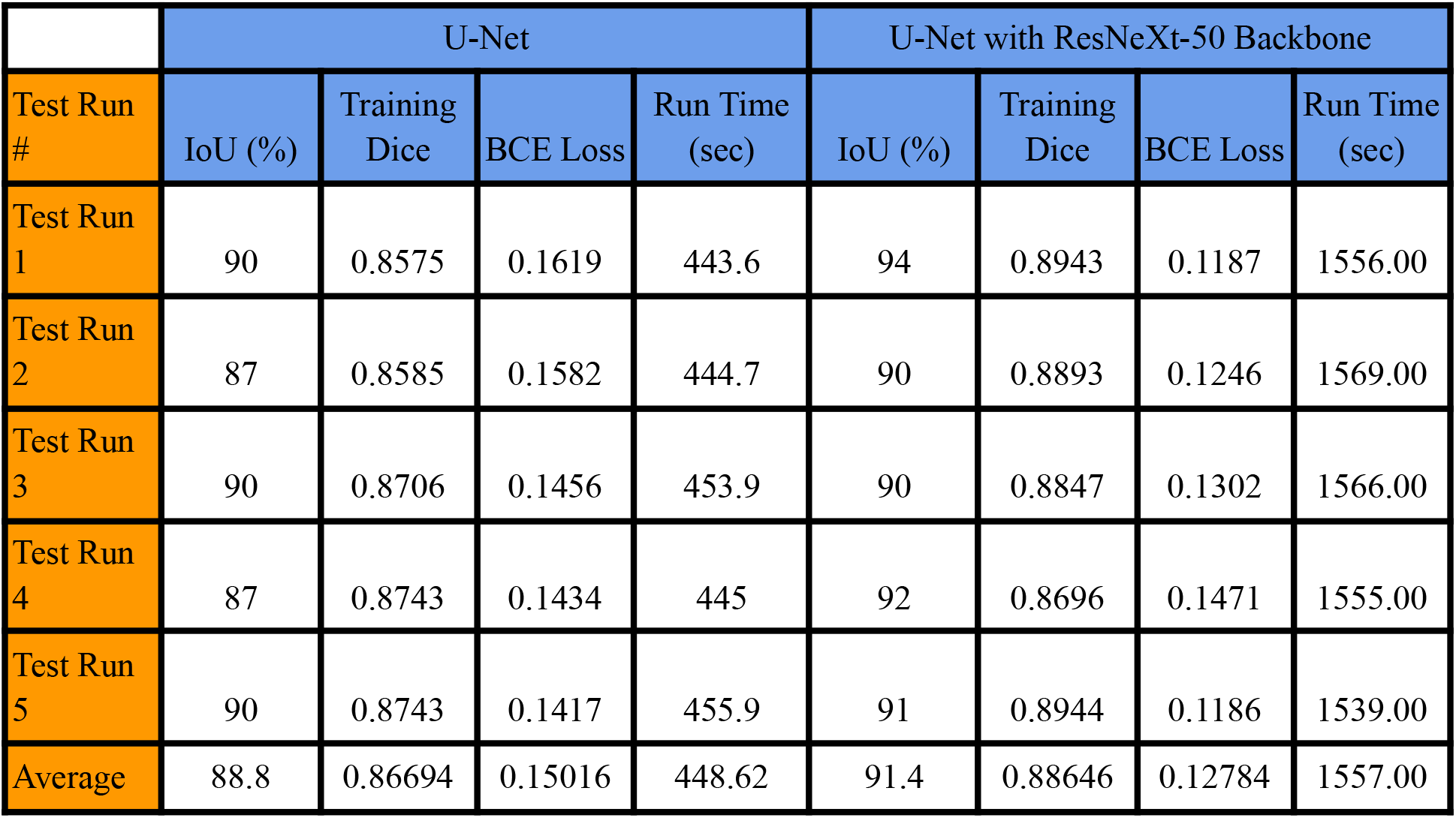
Automatic LGG Segmentation Model Training & Testing Performance

Although both models satisfied the performance criterion for mean training Dice Coefficient (≥ 0.85), the U-Net prototype failed to meet the criteria for mean IoU and BCE Dice loss. The ResNeXt convolutional backbone supported the U-Net architecture and provided higher mean training Dice and mean IOU along with lower mean loss. All performance criteria for segmentation were met through the revised model. Figure 2a and 2b exhibit segmentation predictions from the U-Net with ResNeXt-50 Backbone model.

**Figure 2a.**
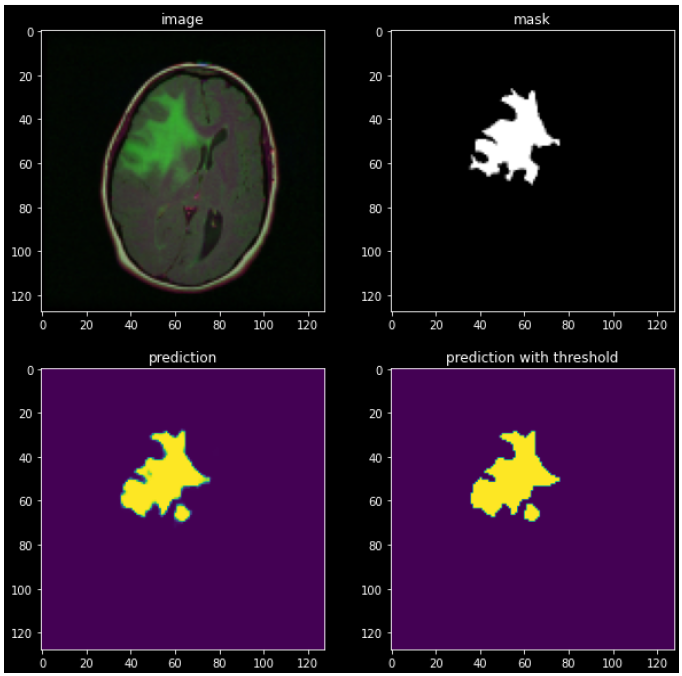
Segmented Tumor Region Prediction of randomly selected LGG MRI

**Figure 2b.**
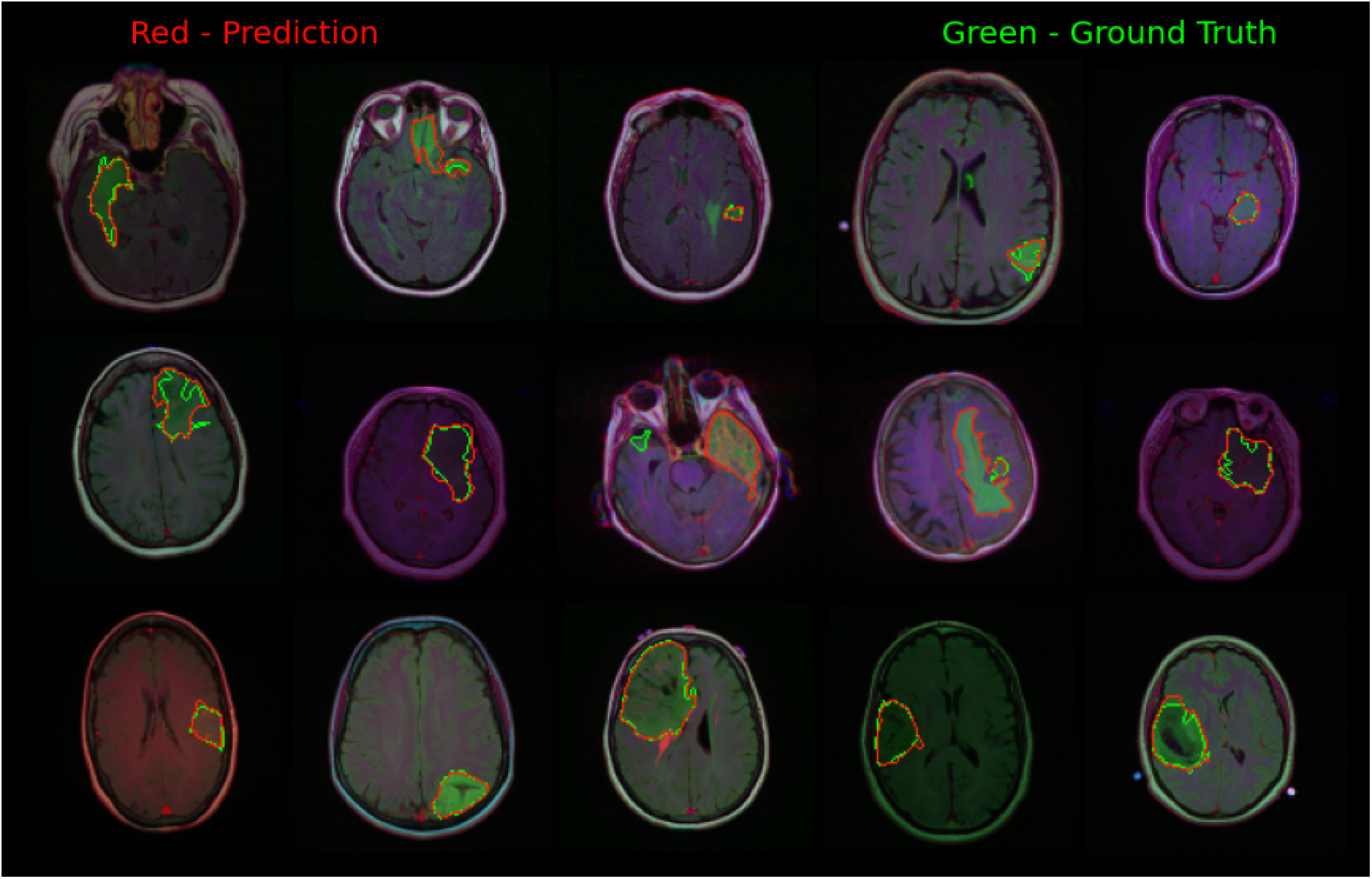
Prediction and Ground Truth Comparison of U-Net with ResNeXt-50 Segmentation Sample

Figure 2a shows and compares the actual tumor FLAIR mask in white and the model’s tumor prediction in yellow. The prediction displayed on the left is based on the basic set of rules defined by the model while the prediction on the right of Figure 2a uses a more stringent threshold to determine tumor regions in MRI. Figure 2b depicts a sample of test image predictions from the redesigned model. It produces a visual comparison between the ground truth FLAIR mask in green and automatic segmentation outlined in red.

### Shape Feature Extraction

Figure 3 visualizes graphs and plots of shape features (MF, and BEVR) for one sample image. Figure 3a shows the calculated centroid of a random tumor image which was used to calculate ASD and MF. Figure 3b depicts MF through a plot of the original radial distances and smoothened radial distances from the tumor centroid to each pixel (represented by pixel number) on the tumor perimeter. Figure 3c illustrates a 3D view of a tumor (in green) surrounded by its minimum bounding ellipsoid (in blue). All distance units are represented in pixels.

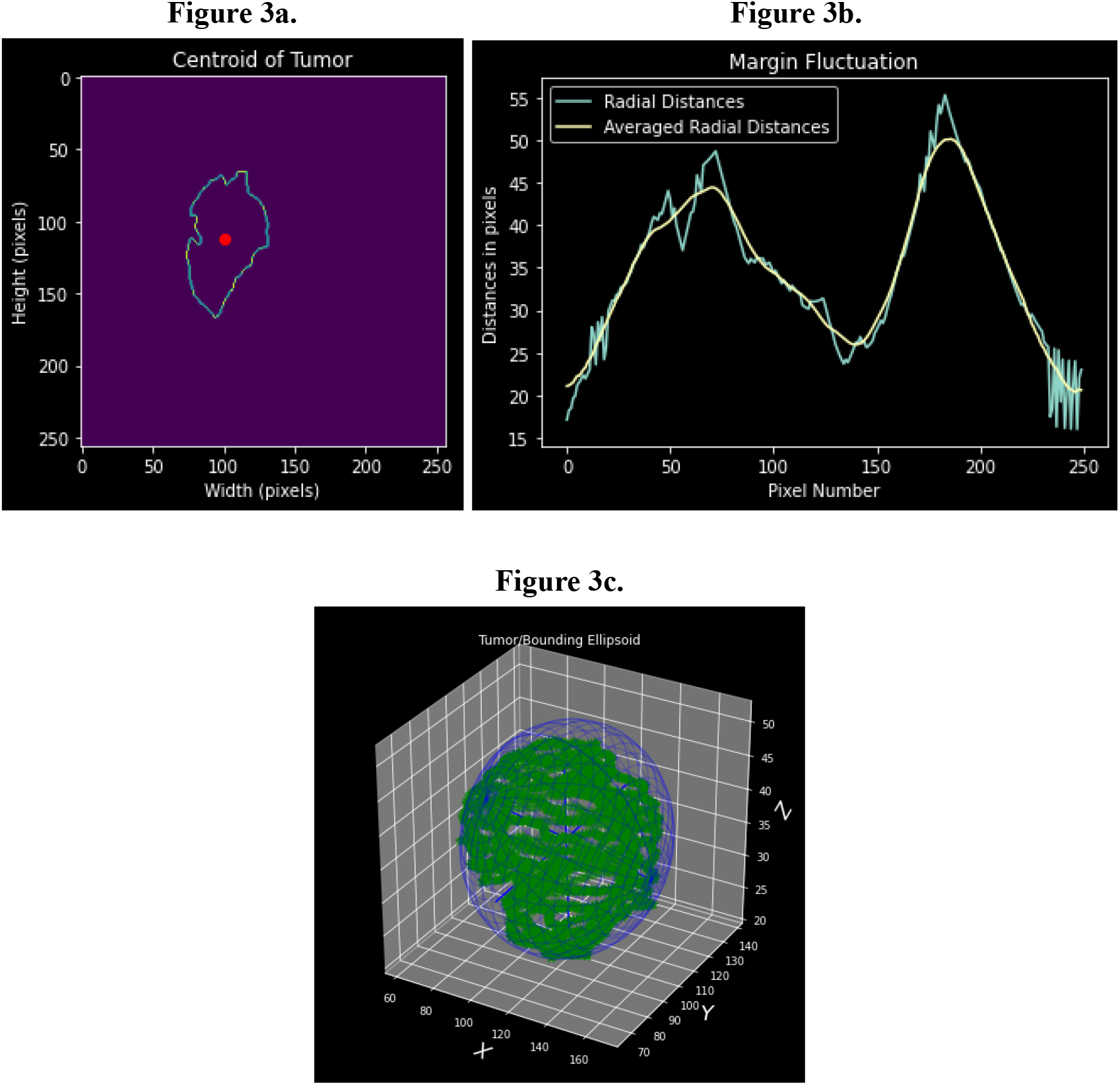

Figure 4 gives the mean values of extracted shape features to quantify tumor characteristics across all 110 patient LGG.

**Figure 4.**
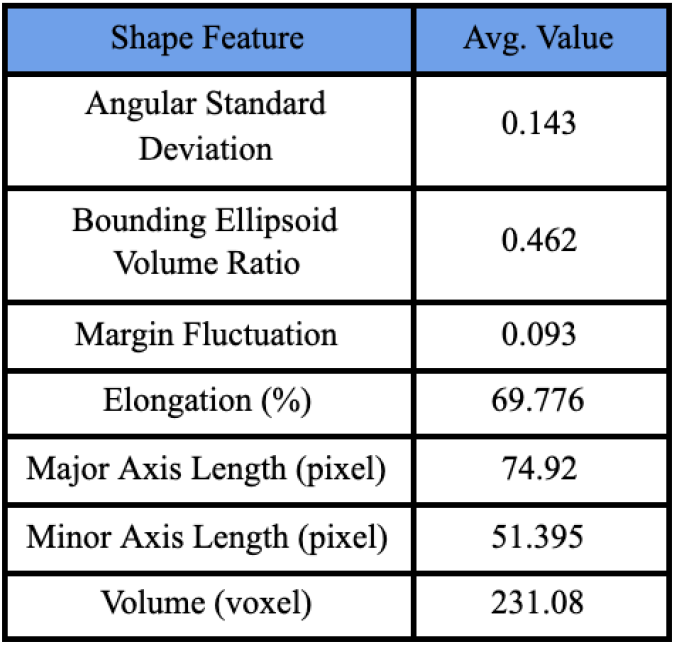
Average Values of Shape Features from Patient LGG Images

An ASD value closer to zero signifies that a tumor is more circular. BEVR values closer to one indicate that tumors have a high degree of 3D regularity. MF values closer to zero indicate that the tumor boundary is relatively smooth and has less high frequency spiculation. Elongation was relatively high, and there was an average of 23.525-pixel difference between major and minor axis length implying that many tumors had high 2D ellipticity.

### Shape Feature - Genomic Subtype Correlation

The final section of the model was using statistical analysis to test relationships or associations between the extracted shape features and provided genomic subtype data. Testing consisted of each shape feature and each subtype (Refer to 2-2 for subtype definitions and 3-2 for shape feature methods) being individually compared through Pearson’s Chi-Squared Tests. Figure 5 shows the results of the 49 tests.

**Figure 5.**
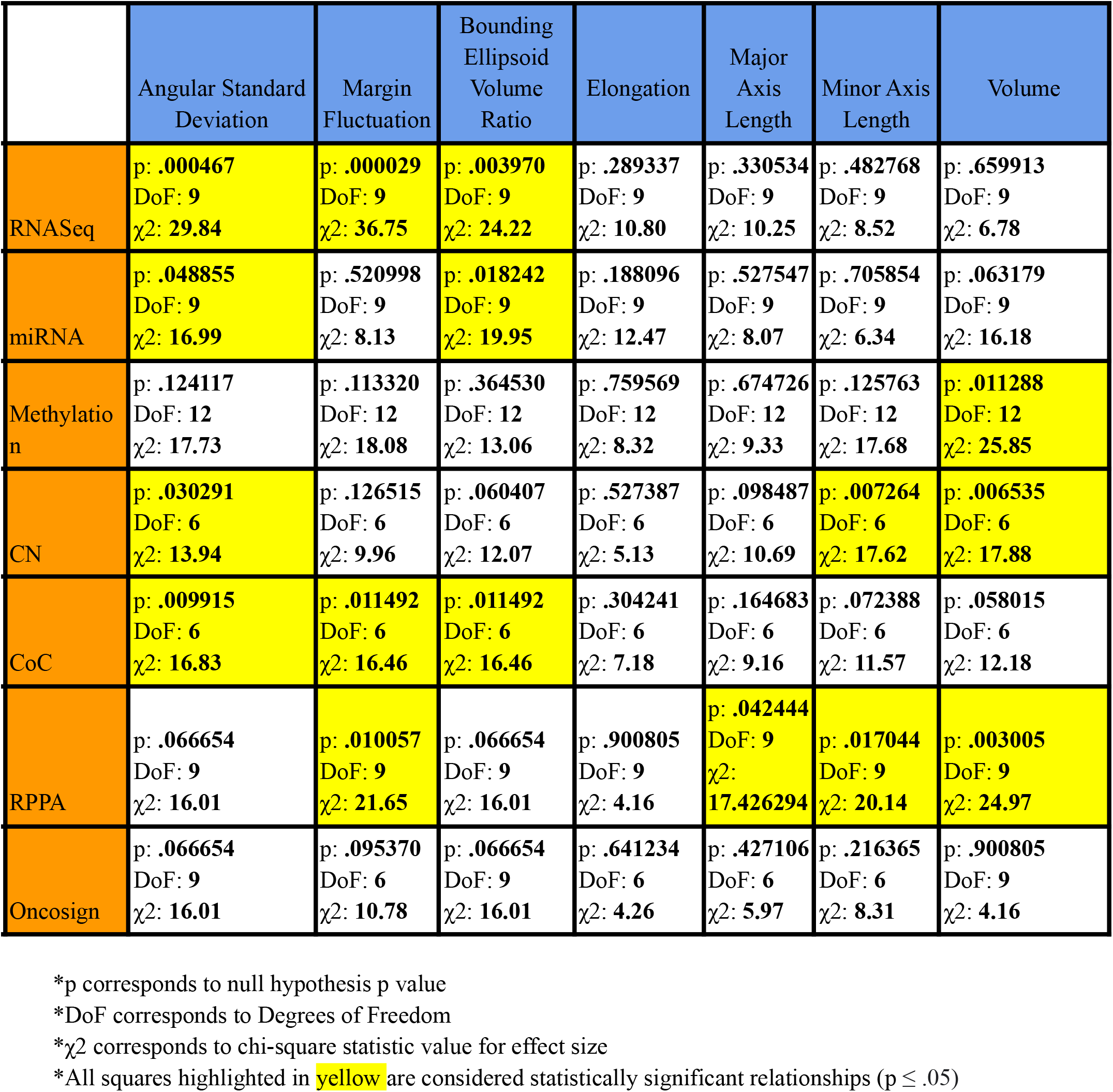
Results of Pearson’s Chi-Squared Tests Between Shape Feature and Genomic Subtype Data

The results of statistical analysis included p-values, degrees of freedom, and chi-square statistic values. P-values in the table indicate the probability of the null hypothesis being true, meaning that no correlation between the tested variables exists. P-values lower than .05 are considered statistically significant as they suggest that the alternative hypothesis, that correlation exists, is at least 95% probable. All highlighted yellow squares had p-values under .05. As can be seen in Figure 5, when p-values were lower, chi-square statistic values were higher, suggesting that lower p-values would have a more sizable effect in the population.

## 5) Limitations and Further Considerations

### 5-1) Limitations

There were limitations of this research project which may have impacted accuracy and results. For segmentation, preprocessing operators were not implemented to execute skull-stripping, which is a denoising step that removes surrounding brain tissue from MRI. This could have altered accuracy. Because 2D images were converted to 3D, shape features calculated using 3D information may have been approximated. Furthermore, shape features were extracted from automatically segmented predictions, so feature values may have been impacted. For statistical analysis, because the patient dataset only provided categorical genomic subtype information, quantitative variable correlation was not possible. Empirical shape feature data had to be categorized by quartile to experimentally test correlation.

### 5-2) Further Research Considerations

- Testing different convolutional backbones in segmentation for increasing accuracy
- Extracting textural tumor features and testing relation with genomic subtypes
- Classification and grading of LGG types based on shape features (astrocytoma, oligodendroma, etc.)

## 6) Conclusion

The use of the reliable convolutional backbone, ResNeXt-50, which supported the U-Net architecture had significant improvement on the performance of the segmentation model. Therefore, the revised model exceeded the performance criteria and provided an effective non-invasive method that can be developed to eliminate the drawbacks of current segmentation efforts. The 91.4% mean testing IoU, 0.9364 mean training Dice, and 0.1237 mean BCE Dice loss that was achieved exceeded several recent models. [7,16] Additionally, the final model required minimal computational expense or runtime. The next step following segmentation was shape feature extraction. This study extracted a combination of 7 novel and previously identified shape features that quantified tumor characteristics. Prior models quantified tumor shape features from imaging datasets by using only the ground truth FLAIR masks. However, one key objective of this model was to create an uninterrupted pipeline that utilizes self-generated segmentation predictions as tumor regions for shape feature extraction. This objective was attained successfully. The shape feature extraction provides useful information about tumors and was used as empirical data for the final section of the project’s framework, correlation testing between shape features and genomic subtypes. In this section it was found, using statistical analysis, that many different shape features and subtypes have potential correlation. The results indicate the following: a) the RNASeq clusters are closely related to ASD, MF, and BEVR, b) the miRNA clusters have potential association with ASD and BEVR, c) Methylation clusters may have correlation with tumor volume, d) CN clusters may have relation to ASD, Minor Axis Length, and Volume, e) CoC clusters could have relation to ASD, MF, and BEVR, and f) there is potential correlation between RPPA clusters and MF, Major and Minor Axis Length, and Volume. Additionally, it was found that smaller p-values are generally associated with larger effect sizes. The relation between MF and RNASeq clusters was the most apparent, as p < .00003 and χ2 = 36.75, indicating that the probability of correlation and the effect size are potentially very high. Oncosign clusters and tumor elongation were not suggestive of any correlation with other groups. Pearson’s chi-squared testing revealed many important results that should be further explored as the existence of correlation could link to better treatment processes for lower-grade gliomas in the future.

## 7) Appendix

### Terms and Formulas

#### IoU

The Intersection over Union (IoU) metric, also referred to as the Jaccard index, is a method used to quantify the percent overlap between the target mask and prediction output.

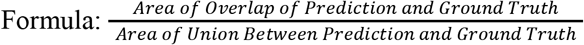

#### Training Dice Coefficient

The Training Dice Coefficient is another method used to quantify the similarity between the target mask and prediction output in each epoch

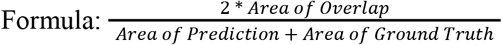

#### BCE Dice Loss Function

The Binary Cross-Entropy (BCE)-Dice Loss Function is a combination of the Dice loss and BCE Loss which measures how much predictions deviate from ground truths after each epoch. The higher the loss, the more variance between model predictions and ground truths.

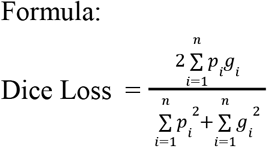

Where p_i_ & g_i_ are corresponding pixel values for ground truth and prediction and N is the total number of pixels in the image.

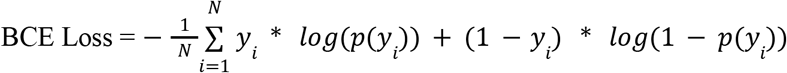

y_i_ is the predicted pixel coordinate at the ith iteration of the series and N is the total number of pixels in the image

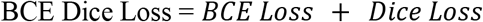

#### Chi-Square Statistic

The chi-square statistic is an extension of Pearson’s chi-squared test which gives a relative effect size of an observed relation on a population

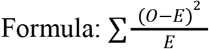

Where O is the observed or actual value and e is the expected value

#### Epoch

The number of epochs is the number of times the training algorithm will iterate through the entire training set

#### Learning Rate

The learning rate is a tuning parameter which defines the rate at which a model corrects its predictions based on weighted observations during training.

#### Batch size

Batch size is the number of objects the training algorithm processes in each sub-epoch iteration.

#### Pearson S Chi-Square Test

Pearson’s Chi-Square Test is a statistical assessment to determine how likely it is that any observed relation between two categorical variables arose by chance

#### P-Value

P-values are determined using the chi-squared test. They are the probability that the null hypothesis, which is the hypothesis that no correlation exists between the evaluated variables, is true. P-values lower than .05 are generally considered as statistically significant because it means that the alternative hypothesis, that a correlation between the observed variables exists, is at least 95% probable.

